# Laminin α1 orchestrates VEGFA functions in the ecosystem of colorectal carcinoma

**DOI:** 10.1101/099465

**Authors:** E. Mammadova-Bach, T. Rupp, C. Spenlé, I. Jivkov, P. Shankaranarayanan, A. Klein, L. Pisarsky, A. Méchine-Neuville, G. Cremel, M. Kedinger, O. De Wever, N. Ambartsumian, S. Robine, E. Pencreach, D. Guenot, J.G. Goetz, P. Simon-Assmann, G. Orend, O. Lefebvre

**Author notes:** equal contribution (T. Rupp, C. Spenlé). co-corresponding authors **Correspondence**: Jacky G.Goetz, Gertraud Orend, Olivier Lefebvre.

## Abstract

Tumor stroma remodeling is a key feature of malignant tumors and can promote cancer progression. Laminins are major constituents of basement membranes that physically separate the epithelium from the underlying stroma. By employing mouse models expressing high and low levels of the laminin α1 chain (LMα1), we highlighted its implication in a tumor-stroma crosstalk, thus leading to increased colon tumor incidence, angiogenesis and tumor growth. The underlying mechanism involves attraction of carcinoma-associated fibroblasts by LMα1, VEGFA expression triggered by the complex integrin α2β1-CXCR4 and binding of VEGFA to LM-111, which in turn promotes angiogenesis, tumor cell survival and proliferation. A gene signature comprising LAMA1, ITGB1, ITGA2, CXCR4 and VEGFA has negative predictive value in colon cancer. Together, this information opens novel opportunities for diagnosis and anti-cancer targeting.

## Introduction

Cancer progression is a multistep process, where the crosstalk between tumor, stromal cells and extracellular matrix (ECM) fosters the survival, proliferation and invasion of tumor cells in (1). Tumor cells and tumor associated stromal cells such as endothelial cells, carcinoma associated fibroblasts (CAFs) and immune cells secrete soluble factors and express specific ECM molecules which are different from that of normal tissues (1).

Laminins (LMs) are heterotrimeric glycoproteins essential for the formation of a highly organized basement membrane (BM), which serves as a barrier between epithelial and mesenchymal tissues (2). The LM family comprises at least 15 described isoforms of LM trimers that are composed of an α, β and γ chain (3). We and others have shown that LMα1 knock out (KO) mice die *in utero* due to the absence of the extra embryonic Reichert’s BM (4,5). However, mice with a Sox2-driven conditional KO of LMα1 (LMα1^cko^) in embryonic tissues or a point mutation in the LN domain (Y265C) of LMα1 are viable (6,7). Yet, these mice exhibit several defects in the retina and the central nervous system (8,9). In humans, patients with biallelic mutations in LAMA1 display cerebellar dysplasia with occasional retinal dystrophy (10,11) altogether suggesting a pivotal role of LMα1 in tissue homeostasis.

As major components of BMs, LMs play a well-known role in cancer, where they could be considered as barriers for cancer cell dissemination. For example, high levels of LMs were found in serum of patients with cancer of the ovary, breast and upper gastrointestinal tract (13–17)

Recently, analysis of tumoral exosomes revealed the presence of LMs (17,18) and high expression of LMs in serum correlates with poor prognosis in colorectal carcinoma patients (15,19). LM-111 has been suggested to promote a malignant phenotype such as enhanced lung metastasis formation from melanoma cells (20). Moreover, tumor cells with forced expression of LMα1 in tumor cells favors tumorigenesis in immune compromised mice (21,22). Yet, very little is known about the underlying mechanisms.

Here, we have used novel transgenic mouse models with ectopic expression of LMα1 in the intestinal epithelium to determine the effects of LMα1 on colon tumorigenesis upon chemically (AOM/DSS) (23) and genetically (mutated APC) (24) triggered tumor induction. We demonstrated that LMα1 promotes tumor formation and angiogenesis by initiating an intimate crosstalk between cancer and stromal cells. We observed that CAFs are attracted by LMα1, and induce VEGFA expression that is tightly regulated by a newly-identified complex of integrin α2β1 and CXCR4. VEGFA binds to LM-111, increasing tumor cell survival and proliferation. Our newly identified signaling axis comprising LMα1, integrin α2β1, CXCR4 and VEGFA correlates with shorter relapse-free survival in colorectal cancer patients. This information may open novel opportunities for diagnosis and targeting of colorectal cancer.

## Results

### Colon tumor incidence and growth is enhanced in transgenic mice overexpressing LMαl

To study how carcinogen treatment affects colon tumorigenesis in context of high LMα1 we have generated transgenic mice with high expression of LMα1. The murine LMα1 cDNA was cloned and expressed under the control of the villin promoter (vLMα1, **Fig. 1A**), driving specific and high expression of the transgene in the epithelium of the gut and along the entire colon (25,26). Resulting transgenic mice (vLMα1) strongly express LMα1 in the colon whereas LMα1 levels were very low in wildtype littermates (**Fig. S1A, B)**. Moreover, LMα1 is expressed in the BM of the colon crypt region of transgenic mice, but not in normal control colon tissue (**Fig. 1B, C**).

**Figure 1.**
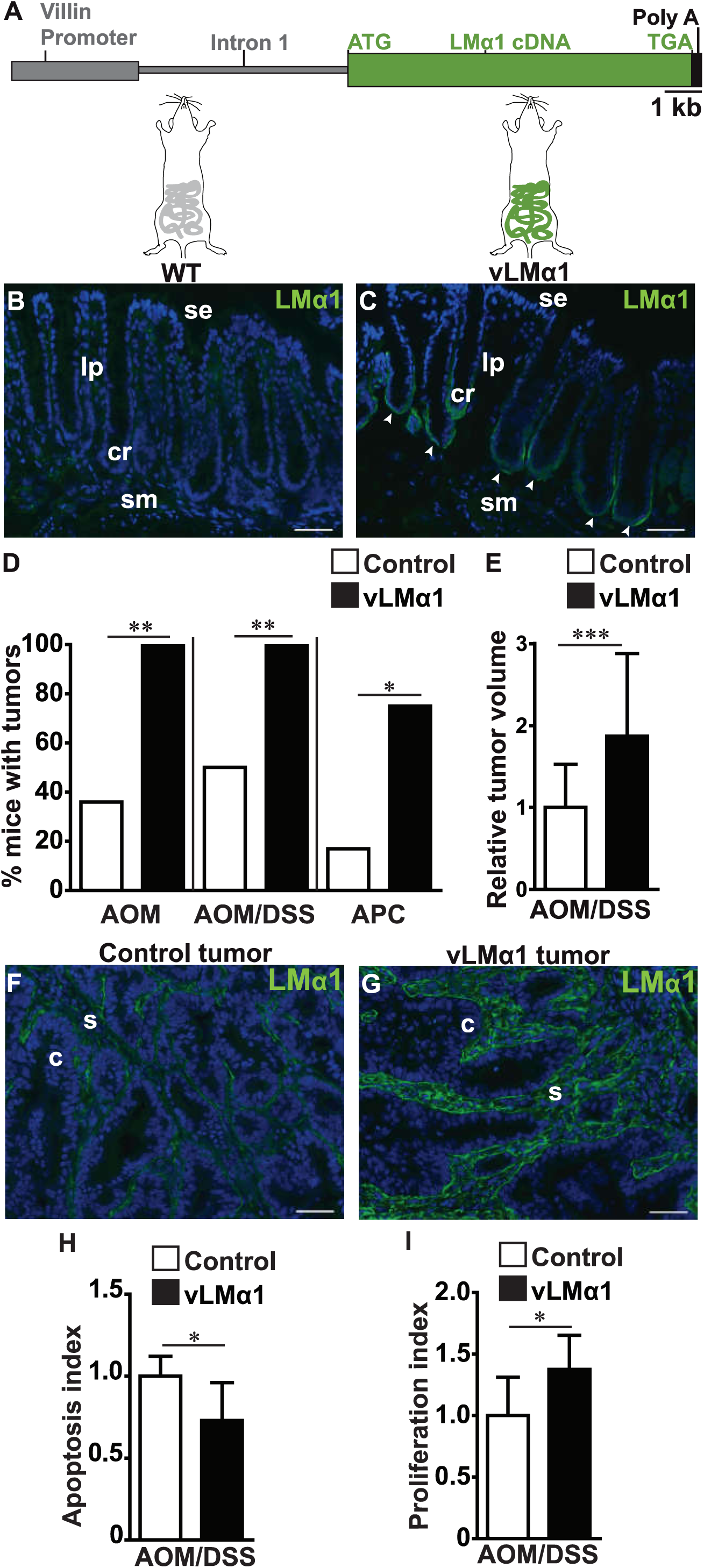
Impact of LMα1 on colon cancer incidence and growth. **A**, Schematic representation of the LMαl transgene. The murine 3.4 kb villin promoter sequence is followed by the 5.5 kb first intron and the 9 kb murine LMal cDNA. **B, C**, LMal tissue distribution assessed by IF with a LMal antibody (green) in tissue from normal distal colon of a wild type (**B**) and vLMαl transgenic mice (**C**). Nuclei were stained with DAPI (blue). Note LMαl expression in the BM of the crypt region (arrowheads), se, surface of epithelium; cr, crypt region; Ip, lamina propria; sm, sub-mucosa. Scale bar: 50pm. **D**, Tumor incidence in control and vLMαl AOM (N=10), AOM/DSS treated (N=12) or APC+/1638 (N=6) mice, chi-square test. **E**, Relative tumor volume in AOM/DSS treated control (n=18), vLMαl (n=68). Fold change, mean+/−SD, permutation test. **F, G**, Tissue expression of LMal assessed by IF with a LMal antibody (green) in colon tumors from AOM/DSS treated control (**F**) and vLMαl (**G**) mice. Nuclei were stained with DAPI (blue), c, cancer cells, s, stroma. Scale bar: 50μπη. **H**, Apoptosis in tumor lysates determined by quantification of activated caspase-3/7. Fold change, mean+/−SD, N=8 mice, n=8 tumors, permutation test. **I**, Proliferation determined by counting Ki67 positive cells per randomly chosen tumor area. Fold change, mean+/-SD, N=8 mice, n=8 tumors, Student’s t-test.

High levels of LMα1 had no impact on animal health or survival and did not induce any macroscopical tissue abnormalities in the gut of older mice. Finally, LMα1 overexpressing mice did not spontaneously develop tumors even after 48 months (data not shown), altogether suggesting that high LMα1 *per se* does not induce tissue abnormalities nor does it foster tumor formation.

Colon carcinogenesis in the mouse was induced by administration of carcinogenic azoxymethan (AOM) or a combination of AOM together with pro-inflammatory dextran sodium sulfate (DSS) in the drinking water (23). Both models lead to intramucosal colon carcinoma formation with different states of transformation after 9 months (AOM) and 2 months (AOM/DSS), respectively eventually leading to a pT2 phenotype (**Fig. S1C, D**). These models closely mimic the human pathology in terms of consecutive events, evolving from *in situ* (pTis) to infiltrating (pT1-pT2) tumors (23,27). To determine whether LMα1 enhances colon tumorigenesis, we assessed the tumor incidence in vLMα1 transgenic mice subjected to carcinogen treatment (**Fig. 1D**). Carcinogen treatment leads to tumors in every single vLMα1 mouse (100%) in contrast to only 36% and 50% of wildtype littermates (AOM,AOM/DSS respectively). When vLMα1 expressing mice were crossed with APC^+/1638N^ mice, that lack an allele of the APC tumor suppressor gene (24), we observed a 75% intestinal tumor incidence in compound vLMα1/APC+^/1638N^ mice, that was higher than the 17% tumor incidence seen in control littermates (**Fig. 1D**). These results suggest that gut specific overexpression of LMα1 significantly promotes tumorigenesis in the chemically and genetically induced carcinogenesis models. We then focused the rest of the study on the AOM/DSS treatment that leads to earlier tumor onset. First, AOM/DSS treated vLMα1 mice have increased tumor numbers and volume (**Fig. 1E**, **S1E**), with tumor displaying increased mRNA and protein expression of LMα1 (**Fig. 1F, G**, **S1F**). LMα1 was strongly expressed in the stroma of AOM/DSS-induced tumors with a high abundance in close vicinity of the cancer cells suggesting that LMα1 may be expressed by tumor and stromal cells (**Fig. 1F**, **G**). We then determined apoptosis and proliferation by measuring caspase 3/7 activity and Ki67 expression, respectively. AOM/DSS treatment reduces apoptosis and increases the proliferation index in vLMα1 mice (**Fig. 1H, I**), suggesting that LMα1 mediates a potential synergism in survival and proliferation during tumor onset and growth. This pivotal role for LMα1 in colon carcinogenesis, was further supported using two grafting models with cells that express abundant and lowered levels of LMα1. Subcutaneous injection of HCT116shLMα1 colon carcinoma cells (**Fig. S1G**) led to smaller tumors than HCT116 control cells which endogenously express LMα1 (**Fig. S1H)**. Similarly, HT29LMα1 tumors overexpressing LMα1 (21) are bigger than HT29 control tumors (**Fig S1I**). Altogether our results showed that LMα1 abundance correlates with tumor incidence and growth in the four analyzed tumor models.

### LMα1 promotes tumor cell survival, proliferation, angiogenesis and pericyte coverage

Angiogenesis and pericyte coverage of vessels are important drivers of tumor growth (28,29). We thus determined vessel density in AOM/DSS induced vLMα1 colon tumors. Increased expression of LMα1 correlated with increased vessel density in mice treated with AOM/DSS (**Fig. 2A-C**). In addition, pericyte-covered blood vessels were increased in carcinogen-induced vLMα1 tumors (**Fig. 2D-F**). Similarly, vessel density and pericyte coverage were increased in both grafted tumor models expressing high LMα1 protein levels (**Fig. 1C, F**, **S2A-H**). In summary, our results demonstrate that LMα1 promotes formation of tumor blood vessels and coverage by pericytes.

**Figure 2.**
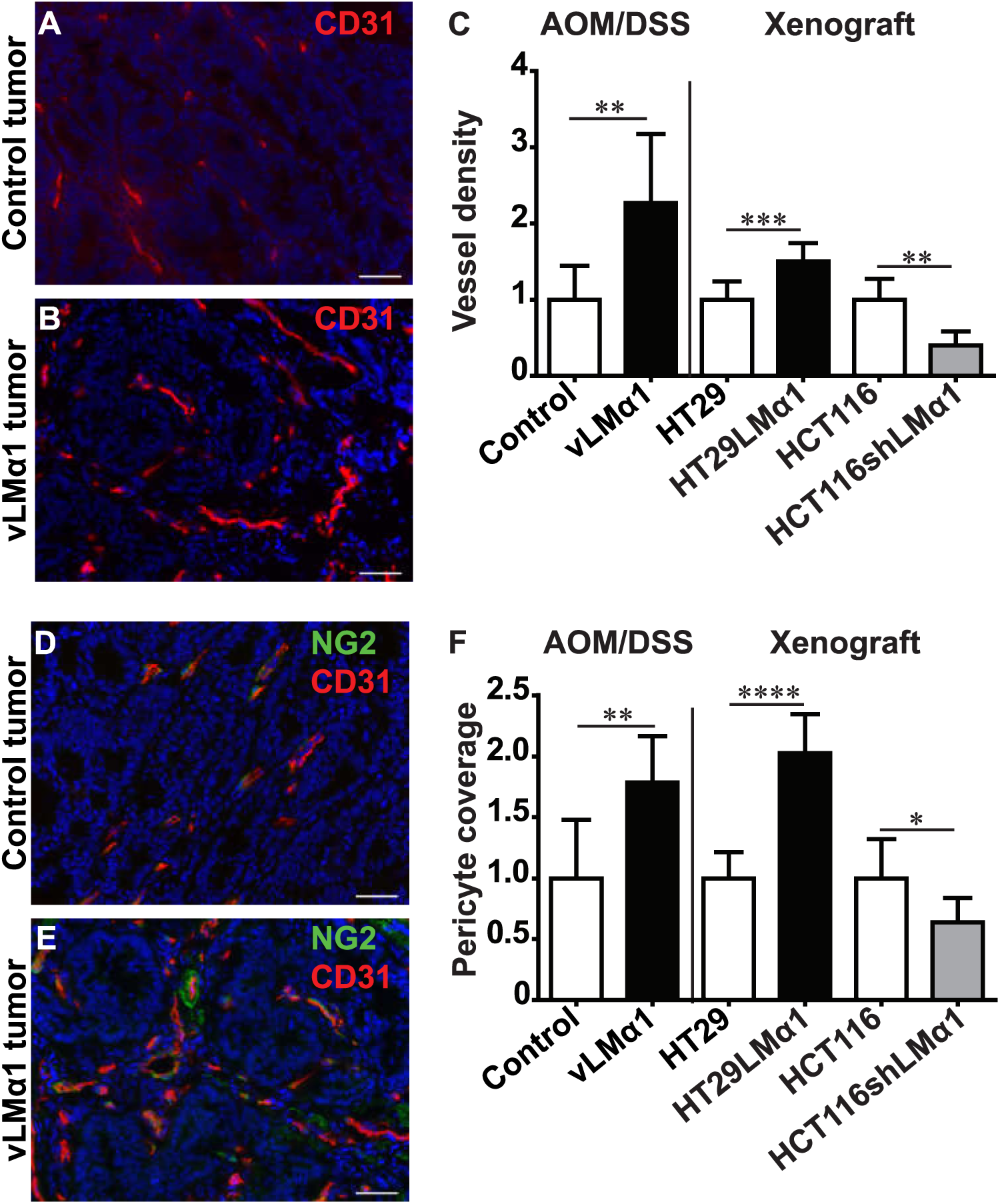
Impact of LMαl on apoptosis, proliferation, tumor angiogenesis and vessel maturation. **A**, **B**, **D**, **E**, Representative examples of IF staining results on sections from AOM/DSS control (**A**, **D**) and vLMαl (**B**, **E**) tumors for CD31 (red, **A**, **B**), combined NG2 (green) and CD31 (red, **D**, **E**). Nuclei were stained with DAPI (blue). Scale bar = 50pm. **C**, Assessment of vessel density by quantification of CD31 signal per area of tumor tissue. Fold change, mean +/−SD, N=8 mice, n=8 tumors, Student's t-test; for HCT116 tumors, permutation test. **F**, Pericyte coverage in AOM/DSS induced colon cancer or xenograft tumors determined as ratio of NG2/CD31 staining per area. Fold change, mean +/−SD, N=8 mice, n=8 tumors, Student’s t-test; for HCT116 tumors, permutation test.

### Carcinoma associated fibroblasts are attracted by LMα1

Since CAFs can stimulate angiogenesis (30) and are increased in tumor xenografts overexpressing LMα1 (21), we assessed the abundance of CAFs in our models. Both staining for αSMA and S100A4, which are common markers for CAFs, were increased in vLMα1 transgenic mice treated with AOM/DSS (**Fig. 3A-F**). This suggests that increased LMα1 recruits CAFs to the stroma of colon carcinoma. A similar result was obtained in xenograft models with tuned levels of LMα1, where expression levels of LMα1 positively correlate with the presence of CAFs (**Fig. S3A-E**).

**Figure 3.**
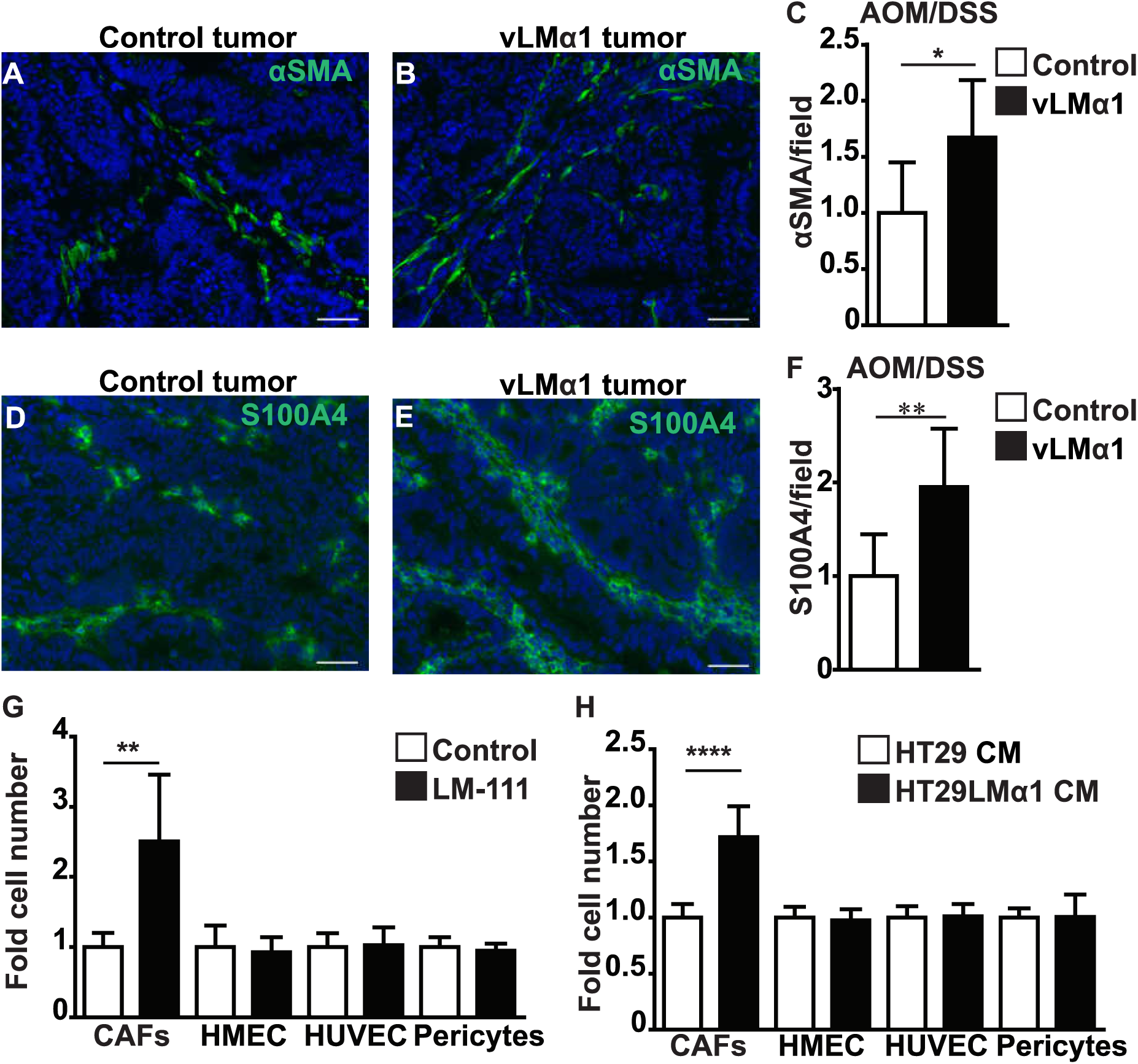
Impact of LMα1 on fibroblast attraction. **A**, **B**, **D**, **E**, Representative images of αSMA (green) positive fibroblasts in AOM/DSS induced control (**A**) and vLMαl (**B**) tumors, of S100A4 (green) positive fibroblasts in AOM/DSS induced colon tumors from control (**D**) and vLMαl (**E**) mice. Nuclei were stained with DAPI (blue). Scale bar = 50pm. **C**, Quantification of αSMA staining signal per area on tissue sections from AOM/DSS induced colon cancer. Fold change, mean+/−SD, N=8 mice, n=8 tumors, permutation test. **F**, Relative S100A4 signal per area on tissue sections from AOM/DSS control and vLMal tumors. Fold change, mean+/−SD, N=8 mice, n=8 tumors, permutation test. **G**, **H**, Boyden chamber assay to assess haptotaxis of CAFs, HMEC, HUVEC and pericytes towards LM-111 or an uncoated surface (control) (**G**) or towards condition medium from HT29LMα1 and control HT29 cells (**H**). Fold change, mean+/-SD, 3 independent experiments in triplicate, permutation test.

The observation that CAFs, endothelial cells and pericytes are recruited in stroma of tumors with higher LMα1 levels prompted us to test the possibility that LMα1 attracted them into the TME. We thus assessed *in vitro* transmigration of CAFs, endothelial cells and pericytes in a Boyden chamber assay towards purified LM-111 (**Fig. 3G**) or conditioned medium (CM) collected from HT29LMα1 cells (**Fig. 3H**). We found that LMα1 enhanced CAF transmigration without stimulating transmigration of endothelial cells (HMECs, HUVECs) or pericytes (**Fig. 3G, H**). This data suggest that while LMα1 efficiently attracts CAFs, but not endothelial cells nor pericytes, which could explain the increased abundance of CAFs in the LMα1-overexpressing tumors.

### LMα1 induces expression of pro-angiogenic molecules

Using a RNAseq approach, we took advantage of the HT29 grafting model that allows to discriminate between stromal and host contribution to LMα1-triggered responses due to assignment to mouse (stroma) and (human) tumor specific genes using the Xenome algorithm (31). We performed genome wide expression analysis of tumors (RNAseq, GEO accession number GSE84296) and cultured cells (Affymetrix, GEO accession number GSE83747). This analysis revealed a significant differential expression of genes in dependence of LMα1 (**Tables S1 to S3**). Gene Ontology analysis and data found in the literature revealed that more than 17% of the genes (62 out of 362) induced in stromal cells, encode molecules regulating angiogenesis (**Fig. 4A**, **Table S4**). In addition, cancer cells also highly express pro-angiogenic genes in the LMα1 overexpressing tumors (**Fig. 4B**, **Table S5**). However, the stromal and tumor pro-angiogenic transcriptomes are mostly distinct. Interestingly, the list of angiogenesis related genes of stromal origin included several well-known pro-angiogenic molecules such as VEGFA and CXCR4 (**Fig. 4A**, **Table S4**), two candidates we considered potentially relevant in LMα1-associated angiogenesis as corroborated below.

**Figure 4.**
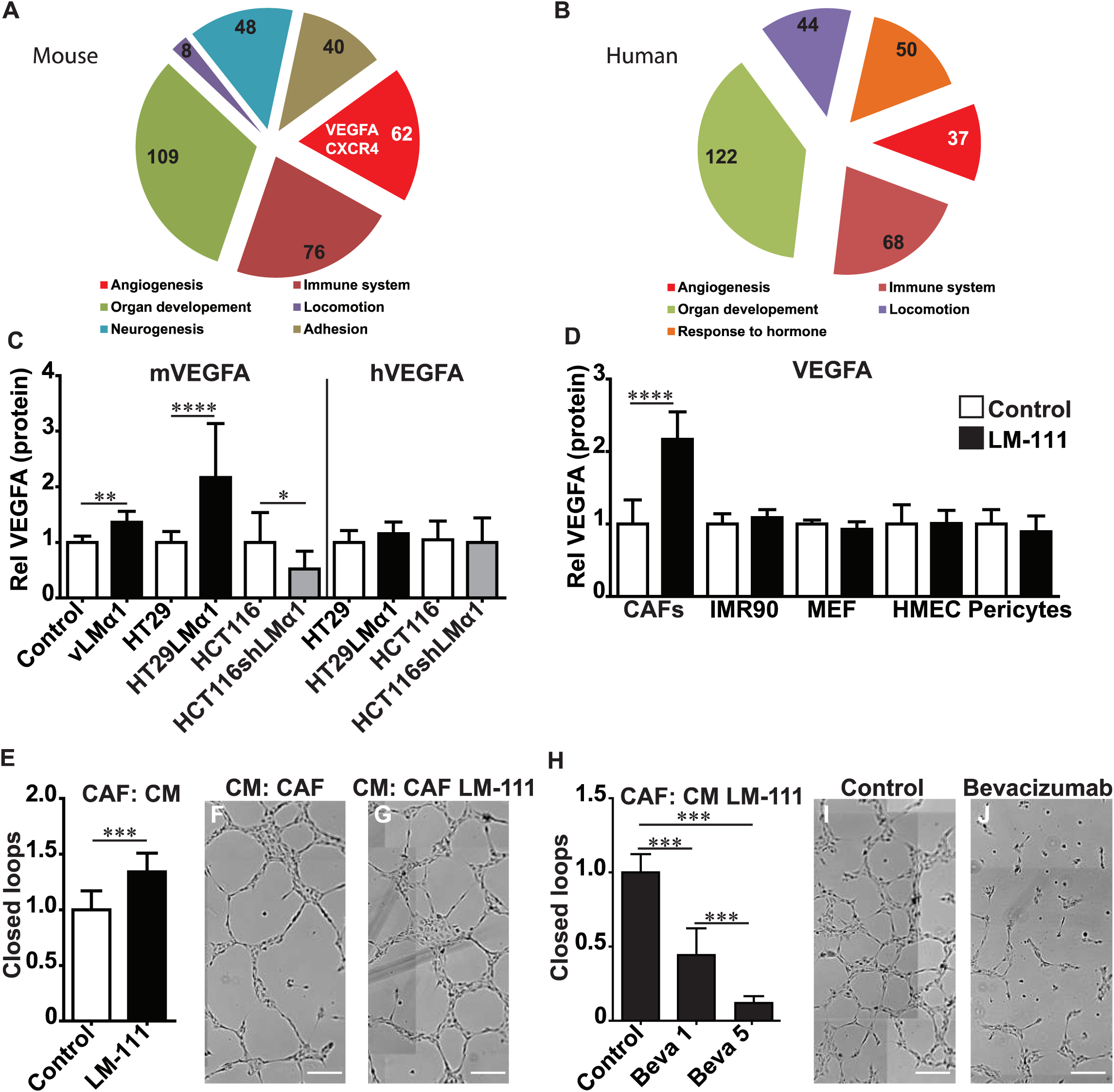
Impact of LMα1 on VEGFA expression and endothelial tubulogenesis. **A**, Pie chart representation of upregulated stromal (mouse) genes implicated in different biological processes and extracted from the RNAseq analysis and using the Panther online tool. Numbers in pie chart are the number of genes in the indicated category among a total of 362 up-regulated genes. **B**, Pie chart representation of upregulated human tumoral genes implicated in different biological processes and extracted from the RNAseq analysis and using the Panther online tool. Numbers in pie chart are the number of genes in the indicated category among a total of 594 up regulated genes. **C**, Species specific expression of VEGFA in AOM/DSS induced colon tumors, and HT29 and HCT116 xenograft tumors was determined by ELISA. Fold change, mean +/−SD, N=8 mice, n=8 tumors, permutation test. **D**, ELISA expression analysis of mVEGFA and hVEGFA (mouse and human respectively) in various cultured stromal cell lines upon cell growth on a LM-111 substratum in comparison to an uncoated surface (control). Fold change, mean+/−SD, 3 independent experiments in triplicate, Student’s t-test. **E**, Quantification of closed loop (tube) formation in HUVEC cells upon exposure to condition medium from CAFs previously grown on a LM-111 substratum or plastic (control). Fold change, mean+/−SD, 4 independent experiments with 5 replicates, Student’s t-test. **F**, **G**, Representative bright field images of HUVEC tube formation upon exposure to conditioned medium (CM) from control CAFs (**F**), CAFs grown on a LM-111 substratum (**G**). Scale bar = 200μιτι. **H**, Quantification of closed loop formation of HUVEC cells upon exposure to condition medium from CAFs previously grown on a LM-111 substratum and treated with the Bevacizumab antibody at 1 (Beva 1) or 5 Mg/mi (Beva 5). Fold change, mean+/−SD, 2 independent experiments with 5 replicates, one way Anova analysis followed by a Tukey post test. **I**, **J**, Representative bright field images of HUVEC tube formation upon exposure to conditioned medium from CAFs grown on a LM-111 substratum, control **(I**) or treated with 5 pg/ml bevacizumab (**J**). Scale bar = 200μm.

### VEGFA expressed by CAFs triggers endothelial tubulogenesis in response to LMα1

We further investigated the role of pro-angiogenic VEGFA in LMα1-associated angiogenesis (32). Moreover, two transcription factors known to induce VEGFA, namely c-Fos (33) and Wt1 (Wilms tumor gene 1, (34) were upregulated 3-fold in the stromal compartment of HT29LMα1 tumors, potentially involved in inducing stromal VEGFA (**Tables S1, S4**). In tumor cells VEGFA levels were not altered irrespective of LMα1 (**Table S1-5**). This was further validated by measuring VEGFA expression in AOM/DSS-induced tumors. We observed increased VEGFA levels following LMα1 upregulation (**Fig. 4C**). Analysis of the tumor xenografts with tuned expression levels for LMα1 further supported this result as VEGFA was found increased in tumors with higher LMα1 levels (**Fig. 4C**, **S4A**). This was not the case for tumor cell derived VEGFA that was unaffected by LMα1 levels (**Fig. 4C**, **S4A**). Altogether, our results suggest that LMα1 elevates VEGFA expression in stromal cells.

We next investigated which stromal cells express VEGFA in response to LMα1 and cultured fibroblasts, endothelial cells and pericytes on a LM-111 substratum before measuring VEGFA expression. We observed increased VEGFA expression only in CAFs and neither in normal fibroblasts (IMR90, MEF), nor in endothelial cells or pericytes (**Fig. 4D**, **S4B**). This observation suggests that LMα1 stimulates VEGFA expression in CAFs, which could act as the primary source for VEGFA-mediated angiogenesis. We thus addressed whether VEGFA contributes to the LMα1-associated angiogenic phenotype. We measured the proliferation and tubulogenesis potential of HUVECs upon addition of conditioned medium (CM) collected from CAFs grown on LM-111. While cell growth was not affected by LM-111 (**Fig. S4C**), CM of LM-111-instructed CAFs increased HUVEC tubulogenesis (**Fig. 4E-G**). We then treated endothelial cells with CM of LM-111-instructed CAFs together with the VEGFA neutralizing antibody bevacizumab (35). This drug treatment largely reduced closed loop formation of the LM-111-instructed CM (**Fig. 4H-J**). These results demonstrate that stromal VEGFA plays a pivotal role in LMα1 promoted endothelial tubulogenesis.

### Laminin-111 induces VEGFA expression in CAFs through an integrin α2β1/CXCR4 complex

We further analyzed how adhesion to LM-111 induces VEGFA expression in CAFs, and focused on integrins and CXCR4. Indeed, integrins mediate adhesion to LM-111 (36,37) and regulate VEGFA expression (38). Additionally, LM-111 increases expression of CXCR4 (39), and CXCL12 and VEGFA can synergistically trigger tumor angiogenesis (40). Blocking integrin α2 or β1, but not α6, significantly reduced VEGFA expression levels in CAFs plated on a LM-111 substratum (**Fig. 5A**, **Fig. S5A)**. Interestingly, only CAFs highly express CXCL12 (**Fig. S5B**) and overexpress its receptor, CXCR4, in response to LM-111 (**Fig. S5C**), validating our RNA-seq analysis of HT29LMα1 tumors indicating elevated expression of CXCR4 in stromal cells (**Table S4**). These results suggest that CXCL12/CXCR4 could mediate VEGFA induction in response to LM-111. We then took advantage of integrin blocking antibodies and observed that blocking integrin α2β1 reduced LM-111 induced CXCR4 expression in CAFs (**Fig. S5D**). We further measured VEGFA expression in CAFs grown on LM-111, upon triggering CXCR4 signaling with CXCL12, and found a further increase in VEGFA (**Fig. 5B**, **S5E**). Moreover VEGFA expression was dependent on integrin α2β1 (**Fig. 5C**, **S5F**). Finally, we observed that LM-111-induced VEGFA expression was effectively CXCR4-dependent (**Fig. 5D**, **S5G**).

**Figure 5.**
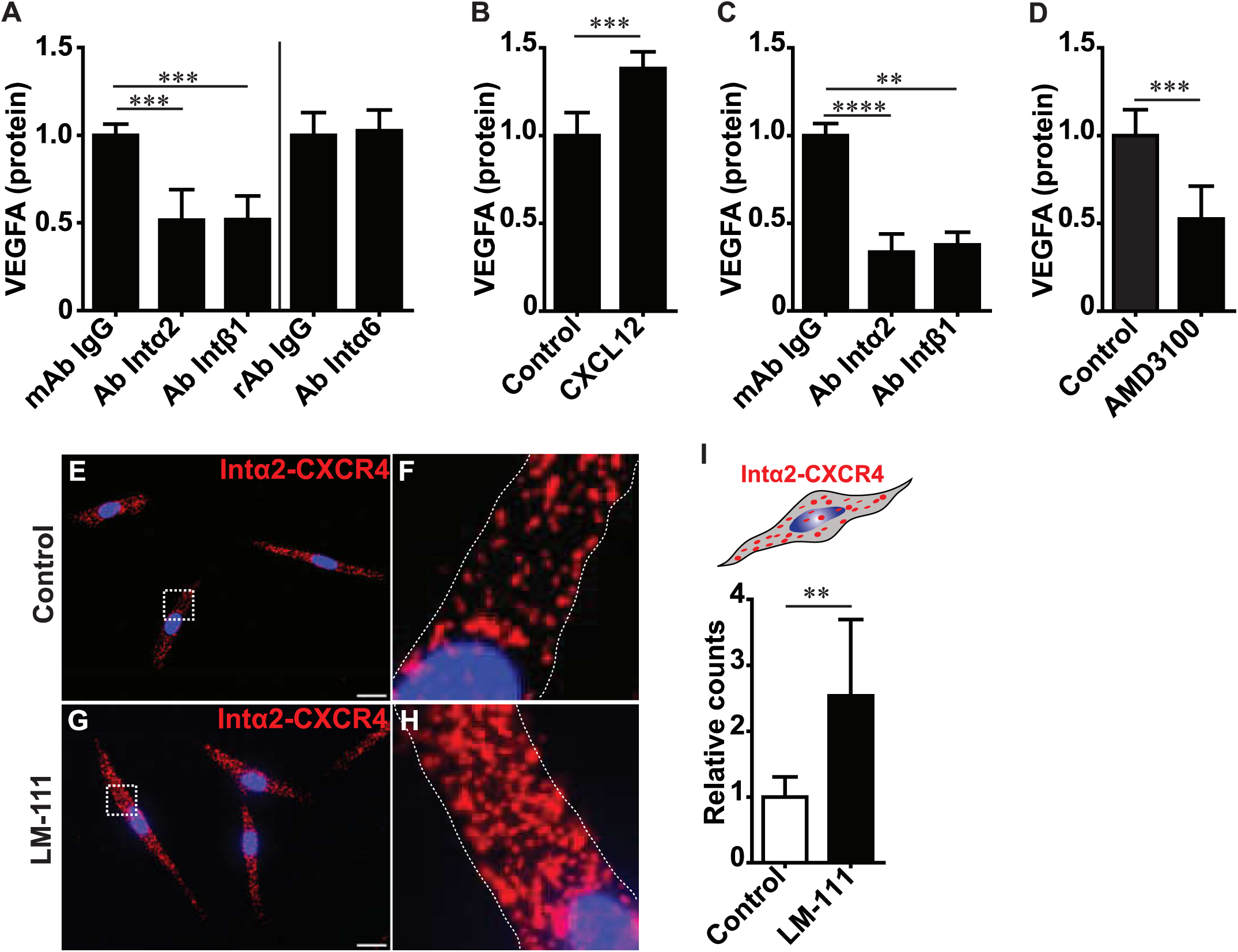
Involvement of α2β1 integrin and CXCR4 in LMα1 triggered VEGFA expression. **A**, Quantification of VEGFA levels (ELISA) in CAFs upon growth on a LM-111 substratum and treatment with control (mouse or rat) IgG or antibodies blocking integrin α2, β1 or α6. Fold change, mean+/−SD, 3 independent experiments in triplicate, one way Anova analysis followed by a Tukey post test. **B**, ELISA expression analysis of VEGFA in CAFs upon growth on a LM-111 substratum and treated with solvent (control) orCXCL12. Fold change, mean+/- SD, 3 independent experiments in triplicate, permutation test. **C**, ELISA expression analysis of VEGFA in CAFs upon growth on a LM-111 substratum, treated with CXCL12 and with control or blocking antibodies against integrin a2 or β1. Fold change, mean+/−SD, 3 independent experiments in triplicate, one way Anova analysis followed by a permutation test. **D**, ELISA expression analysis of VEGFA in CAFs upon growth on a LM-111 substratum and treated with vehicle (control) or the CXCR4 inhibitor AMD3100. Fold change, mean+/− SD, 3 independent experiments in triplicate, Student’s t-test. **E-H**, Representative IF pictures of proximity ligation assay results to visualize the interaction of integrin α2 with CXCR4 in CAFs upon growth on plastic (control, **E**) or a LM-111 substratum (**G**). Respective high magnification pictures are represented in (**F**) and (**H**). Nuclei were stained with DAPI (blue). Scale bar = 20μm. **I**, Proximity ligation assay quantification of integrin α2/CXCR4 complex formation. Mean +/−SD, 2 independent experiments in pentaplicate, Student’s t-test.

Because LM-111 triggered VEGFA expression in CAFs by a mechanism that involves integrin α2β1 and CXCR4 and since inhibition of each molecule blocked LM-111-induced VEGFA expression (**Fig. 5A-D**), we considered a potential interdependence of signaling by a physical complex of the two receptors. We thus exposed CAFs to a LM-111 substratum and assessed interaction of integrin α2β1 and CXCR4 using a proximity ligation assay. While CXCR4 and integrin α2β1 physically interact in steady-state conditions, LM-111 further promotes this complex formation (**Fig. 5E-I**). Altogether, our results suggest that *in vitro*, high levels of tumor cell-derived LMα1 trigger VEGFA expression in CAFs by a mechanism that involves LM-111-dependent integrin α2β1/CXCR4 complex formation.

### LM-111 promotes tumor cell survival and proliferation via its binding to VEGFA

Because CM from LMα1-exposed CAFs promoted endothelial tubulogenesis, we asked whether this CM had also an effect on survival and/or proliferation of tumor cells. Here, we used the HT29 carcinoma cells with tuned LMα1 expression levels. Upon plating HT29 cells on LM-111 together with apoptosis-inducing staurosporine, we observed no effect on apoptosis in the parental cells with CM from CAFs grown on LM-111 or plastic. Surprisingly, less apoptosis was noted when CM from LM-111-instructed CAFs was added to HT29LMα1 cells (**Fig. 6A**). In addition, CM from LM-111-instructed CAFs increased proliferation of HT29LMα1 cells. In particular, an increased number of HT29LMα1 cells entered the S and G2/M phases, which were accompanied by a decrease in the G0/G1 phase (**Fig. 6B**). These data suggest that LM-111-instructed CM of CAFs promotes survival and proliferation of LMα1 overexpressing tumor cells. Thus, LMα1 may promote survival and growth signaling in tumor cells by a synergy between pericellular LMα1 and LMα1-induced soluble factors.

**Figure 6.**
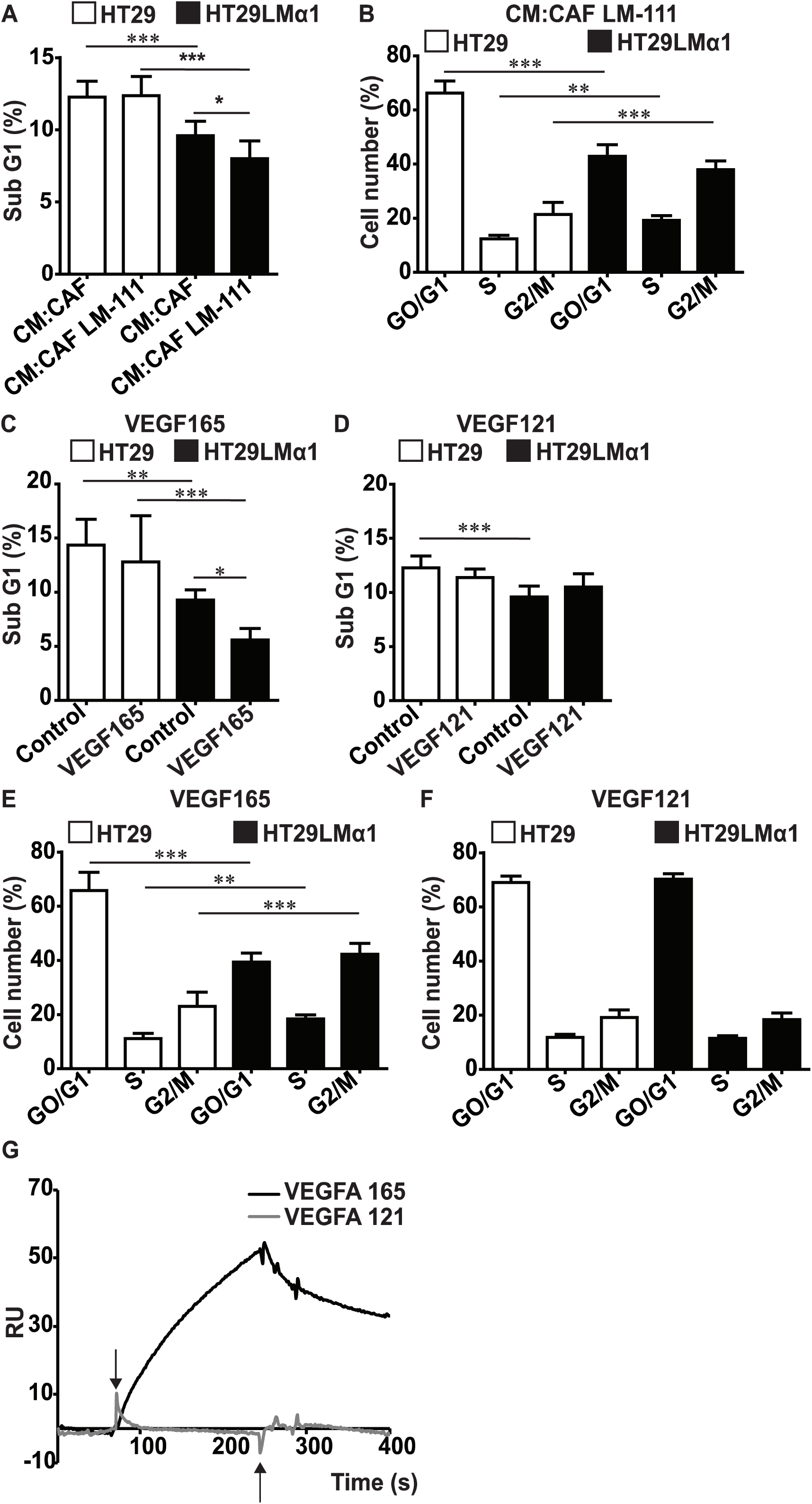
Impact of LMcri/LM-111 on VEGFA binding, survival and proliferation. **A**, FACS analysis of the sub G1 fraction of staurosporin triggered HT29 control and HT29LMα1 cells treated with conditioned medium (CM) from CAFs previously grown on a LM-111 substratum or plastic. Mean+/×SD, 3 independent experiments in triplicate, one way Anova analysis followed by a Tukey post test. **B**, FACS analysis of G0/G1, S, G2/M fractions of HT29 control and HT29LMα1 cells treated with conditioned medium from CAFs previously grown on a LM-111 substratum. Mean+/−SD, 3 independent experiments in triplicate, one way Anova analysis followed by a Tukey post test. **C**, **D**, FACS determination of the sub G1 fraction of staurosporin triggered HT29 control and HT29LMα1 cells treated with recombinant VEGFA165 (**C)** and VEGFA121 (**D**). Mean+/−SD, 3 independent experiments in triplicate, one way Anova analysis followed by a Tukey post test. **E**, **F**, FACS analysis of G0/G1, S, G2/M fractions of HT29 control and HT29LMα1 cells treated with recombinant VEGFA165 (**E**) and VEGFA121 (**F**). Mean+/−SD, 3 independent experiments in triplicate, one way Anova analysis followed by a Tukey post test. **G**, Binding of VEGFA165 or VEGFA121 to LM-111 (adsorbed to a CM5 sensorchip) was determined by Biacore with normalization towards a blank surface. The arrows depict the beginning of the association and dissociation phases respectively (start and end point of injection). Note, contrary to VEGFA121, VEGFA165 bound to LM-111 in a dose dependent manner with a Kd of 4.7}1 O’^8^ M.

As VEGFA largely mimicked the LMα1 effect on endothelial tubulogenesis (**Fig. 4E-J**) and VEGFA had previously been shown to promote tumor cell survival (32), we considered VEGFA as candidate factor of the LM-111-instructed CM promoting survival and proliferation in tumor cells. VEGF121 and VEGF165 isoforms of VEGFA have been shown to display distinct angiogenic properties where VEGFA165 is more active, via its interaction with the co-receptor NRP1 (41). Interestingly, whereas staurosporin-triggered apoptosis was reduced by VEGFA165 in HT29LMα1 cells (yet not in the parental cells), this was not the case with VEGFA121 (**Fig. 6C, D**). Moreover, VEGF165 enhanced entry of HT29LMα1 cells into S and G2/M phases which was accompanied by a reduction in the G0/G1-phase (**Fig. 6E**). No effect was seen when using VEGFA121 (**Fig. 6F**). In summary, our experiments revealed that VEGF165 is promoting survival and proliferation of tumor cells thus mimicking the LMα1 effect. Moreover, response to VEGFA depends on cell adhesion to LMα1 suggesting a combined signaling by LMα1 and VEGFA.

Because LMs and VEGFA possess heparin-binding domains and heparin binding sites respectively (42,43), we considered that VEGFA and LMα1 act via physical interaction. We used surface plasmon resonance spectroscopy to measure binding of VEGFA165 to sensor chip-adsorbed LM-111. We observed that VEGFA165 binds LM-111 in a dose dependent manner with a Kd of 4.7 × 10^−8^ M (**Fig. 6G**), in the range of the previously reported binding of VEGFA165 to heparan sulfated glycosaminoglycans (2.4 × 10^−8^ M, (44). VEGF121 lacks the heparin binding sites (41) and did not interact with LM-111 (**Fig. 6G**). Thus, the heparin binding activity in VEGFA165 is crucial for the interaction with LM-111 to mediate the survival and growth promoting effect of LMα1.

### Prognostic value of the LMα1-integrin α2β1-CXCR4-VEGFA axis for poor survival

Altogether, our results suggest that LMα1 is involved in colon carcinogenesis. We investigated whether this was specific to LMα1 and determined relative expression levels of the twelve LM chains in 39 primary human colorectal cancer specimens (**Table S6**). While LMα5 transcripts were the most abundant (not shown), LMα1 was the most highly induced (18-fold) LM molecule in tumor tissue compared to adjacent non-tumor tissue (**Fig. 7A**).

**Figure 7.**
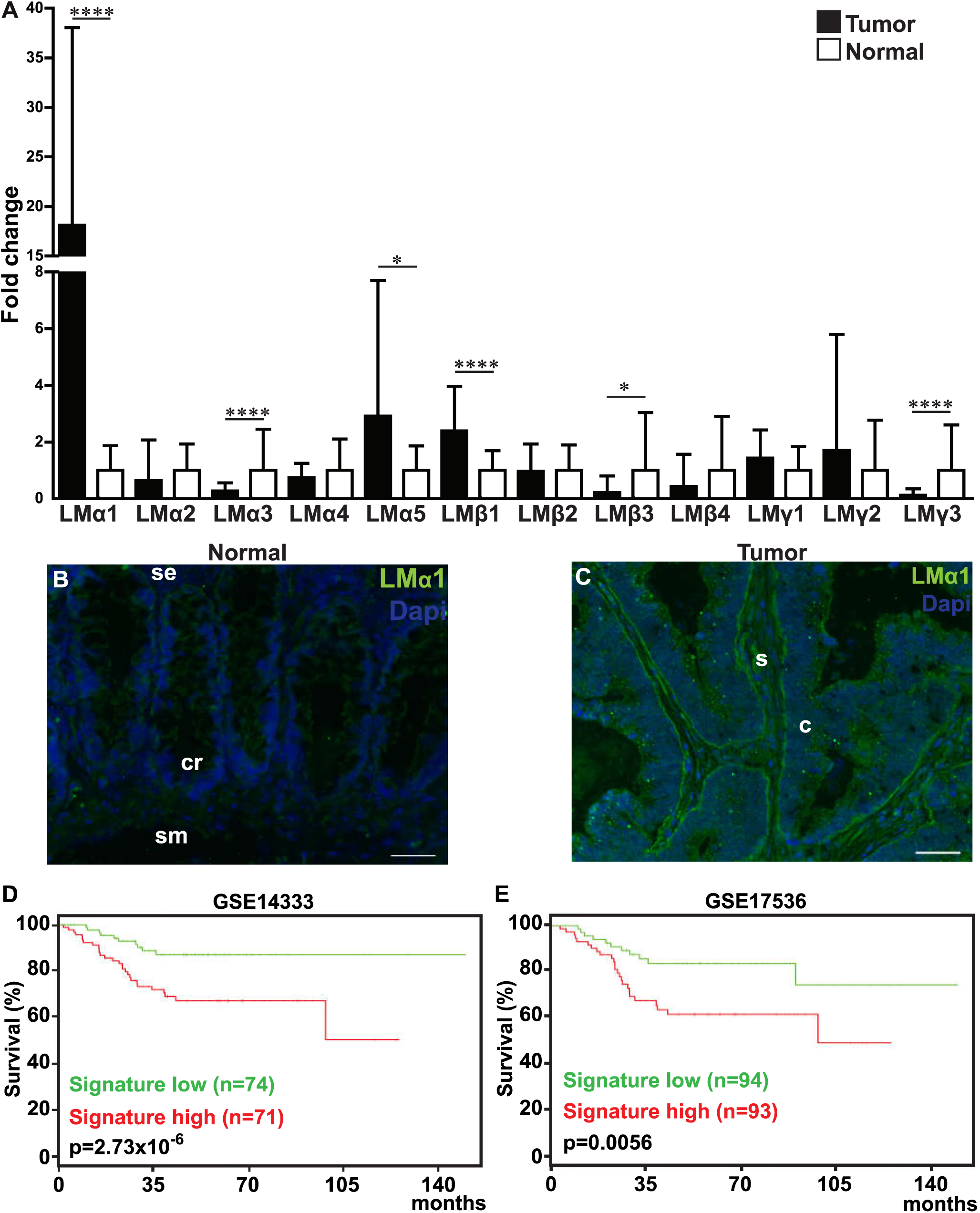
LMα1 expression in human colorectal cancer. **A**, Expression of the twelve laminin chains in human colorectal tumors was determined by qRT-PCR in 39 tumoral tissues and matched normal colon tissues. Fold change, mean+/− SD, permutation test. **B**, **C**, Representative immunofluorescence pictures of LMcrt (green) in normal human colon (**B**) and human colorectal tumor tissue (**C**). Nuclei were stained with DAPI (blue), cr, crypt; se, surface of epithelium; sm, sub-mucosa; c, cancer cells; s, stroma. Scale bar: 50μm. **D**, **E**, Kaplan-Meier survival analysis of colorectal tumor patient cohort GSE14333 (**D**) and GSE17536 (**E**). Data were stratified according to the average expression of LMαl, integrin α2, integrin β1, CXCR4 and VEGFA as high or low using a combined median cutoff value. Note, that in both cohorts high expression of the LMαl associated gene signature significantly correlates with decreased survival, n indicates the number of samples per group, p-values indicate the significance of survival difference between the groups of individuals.

LMα1 mRNA is already highly induced at early tumor stage (pT1) and LMα1 expression levels do not correlate with tumor stage (**Table S6**). In addition to LMα1, transcripts of two other LM chains, LMα5 (2.9-fold) and LMβ1 (2.4-fold) were also significantly overexpressed in colon tumor tissue. In contrast, transcript expression of LMα3 (-3.6-fold), LMβ3 (-4.7-fold) and LMγ3 (-8.4-fold) appeared lower in the tumor tissue (**Fig. 7A**).

We further observed that whereas LMα1 was mostly absent from non-tumor tissue, LMα1 could be found highly expressed in the TME, predominantly at the interface between cancer cells and in the tumor stroma (**Fig. 7B, C**), resembling LMα1 expression in AOM/DSS induced murine tumors (**Fig. 1F, G**). This analysis revealed a complex deregulated expression of LMs in colon tumors, where LMα1 is the most significantly deregulated LM chain.

Finally, we wondered whether stromal LMα1 associated short signature comprising LAMA1, ITGB1, ITGA2, CXCR4 and VEGFA potentially correlates with patient prognosis by using publicly available mRNA expression data sets from two cohorts of patients (n=332) with colorectal cancer (GSE14333 and GSE17536). We observed that combined high expression of the five genes strongly correlates with reduced disease free survival in both cohorts (**Fig. 7D, E**). Our results suggest that LMα1 together with VEGFA, ITGB1, ITGA2 and CXCR4 are interesting markers of patient outcome in colorectal adenocarcinoma with targeting opportunities.

## Discussion

Extracellular matrix, and in particular LMα1, can promote tumorigenesis (21,22) yet the underlying mechanism was elusive. Here, we employed a multi-scale approach *(in vitro* assays, murine models and cancer patients) to discover the molecular mechanism of LMα1 promoted tumor growth and prognostic value for cancer patients (see summary, **Fig.8**).

Overexpression of LMs is not necessarily enhancing the barrier function of BM, that is often lost during cancer progression (22). Indeed some overexpressed LM isoforms were found to promote growth, protease production and angiogenesis (20). We had recently shown that LMα1 is induced in inflammatory bowel disease (IBD) by a mechanism that involves p53. We further demonstrated that transgenic mice overexpressing LMα1 in the intestinal epithelium are less prone to DSS-induced inflammation, suggesting a barrier function of LMα1. Yet, surprisingly the carcinogen AOM induced more intestinal tumor lesions in vLMα1 mice (26). Here, we addressed the mechanisms underlying LMα1 induced tumorigenesis in more details by employing our novel transgenic mouse models expressing high LMα1 levels and a cancer driving mutation in APC (APC^1638N/+^;(24) or by using conditions driving tumorigenesis (AOM or AOM/DSS) that are relevant for the human disease (23). In addition, we used two colon cancer xenograft models with LMα1 overexpression or shRNA-mediated LMα1 knockdown. In all models we observed that LMα1 overexpression enhanced tumor incidence, tumor growth and, most importantly, stromal activation that increased tumor angiogenesis. Our models are relevant for the human disease since we observed a strong induction of LMα1 in human colorectal cancers. In addition, LMα1 is highly abundant in the tumor tissue of colorectal cancer specimens, reaching more than 18-fold increased mRNA levels than in non-tumorigenic tissue.

We further demonstrated that LMα1 orchestrates the TME by triggering a crosstalk between cancer and stromal cells that tunes VEGFA expression. We identified CAFs as central stromal player that are more abundant in LMα1 overexpressing tumors suggesting that LMα1 attracts CAFs into the tumor tissue. Indeed, in cell culture CAFs were attracted by LM-111 which did not apply to other stromal cells.

In addition to VEGFA, CAFs also express CXCL12 that activates CXCR4. Although cancer cells express different chemokine-receptors, CXCR4 is correlated with poor prognosis for colorectal cancer patients (45) and is commonly found in multiple types of cancer, thus strengthening the importance of the CXCL12/CXCR4 axis in progression and metastasis of colon cancer (46). Previously published data suggest a role of CXCR4 on LM-111 dependent tumor cell adhesion and chemotaxis potentially through overexpression of α5 and β3 integrins (47). Here, we demonstrated for the first time that on a LM-111 substratum a physical complex of CXCR4 and α2β1 integrin is stimulated, as shown by a proximity ligation assay. Signaling by LM-111 through integrin α2β1 is crucial for VEGFA induction, as this was blocked by an integrin inhibitory antibody as well as by the CXCR4 inhibitor AMD3100. CXCL12 dependent CXCR4 activation was shown to potentially enhance β1 and in particular α6β1 integrin dependent adhesion of small cell lung and pancreatic cancer cells on LM, respectively (39,48). Although a physical interaction of CXCR4 with integrin β1 may be inferred from a recent publication demonstrating clustering of β1 integrins in CAFs upon stimulation with CXCL12 (49), a physical interaction and dependence on LM-111 has not yet been demonstrated.

We also provide for the first time evidence that LM-111 binds VEGFA165, that occurred with a Kd in the range described for VEGFA binding to glycosaminoglycans (44). The interaction presumably occurs through the heparin binding site in the LMα1 G domain (42,43). In contrast, due to its lack of heparin binding sites (50), VEGFA121 does not bind to LM-111 and thus has no effect on colon tumor cell proliferation and survival. We further demonstrate that LM-111 binds VEGFA165 and that this form is biologically active in promoting colon cancer cell survival and proliferation on LM-111. The latter could also apply to blood vessel formation since VEGFA165 is inducing angiogenesis (41). It is tempting to speculate that binding of VEGFA to LMα1 may play a similar role as binding of VEGFA to NRP1, since both molecules bind VEGFA 165 via their heparin binding sites (51) thus potentially stabilizing VEGFA/VEGFR recognition. Most importantly, the LMα1-integrin α2β1-CXCR4-VEGFA axis correlates with shorter relapse free survival in colorectal carcinoma patients. Thus, the identified crosstalk initiated by high LMα1 expression in cancer cells and involving integrin α2β1, CXCR4 and VEGFA is clinically relevant. This knowledge might be useful for diagnosis and future combination therapies. FDA-approved drugs targeting CXCR4 and VEGFA are currently tested in clinical combination trials in glioblastoma (NCT01339039). Our results suggest that this combination treatment might also be useful for patients diagnosed with colon cancer irrespective of tumor stage since LMα1 is already upregulated early in colon tumorigenesis (shown here) as well as in IBD where p53 induces LMα1 (26). Finally, our novel murine tumor models may be useful in increasing our understanding of vascular BM assembly and barrier function in cancer.

In summary, we demonstrated that ectopically expressed LMα1 in intestinal tissue plays a Janus role. Whereas ectopically expressed LMα1 protects from chronic inflammation induced tissue damage in IBD (26), the present study shows that LMα1 promotes colon tumor incidence, growth and angiogenesis by orchestrating an intricate crosstalk between cells and the LM-111 matrix. LMα1 expressed by colon cancer cells recruits CAFs that secrete VEGFA in response to concomitant signaling by CXCR4 and integrin α2β1 upon adhesion to the LM-111 substratum. In turn, VEGFA binds to the LM-111 rich substratum, which promotes tumor cell survival and proliferation. VEGFA signaling also promotes angiogenesis leading to more and presumably functional vessels thus supporting tumor growth (**Fig. S6**). The identification of VEGFA, CXCR4 and α2β1 integrin downstream of LMα1 in colon cancer is of bad prognostic value for patient survival and may open novel opportunities for targeting colon cancer.

## Material and methods

### Human colorectal cancer specimens

Primary human colorectal tumors of different differentiation state, and normal tissue with no signs of tumorigenesis were obtained from 39 patients with their written consent. All surgical specimens were evaluated and histologically analyzed by an experienced pathologist and classified according to the TNM staging system denominating the tumor (T) stage, presence of lymph nodes (N) and metastasis status (M). Replication error (RER) status of all surgical specimens was determined and provided by the Centre de Ressources Biologiques, (Hôpitaux Universitaires de Strasbourg, Hôpital de Hautepierre, Strasbourg, France). Patient information is listed in Supplementary Table S6. Tumor material and healthy tissue were immediately snap frozen for RNA preparation or was embedded in Tissue-Tek (Labonord, Templemars, France) for immunofluorescence analysis.

### Cloning of the villin-LMα1 vector

Full details of the cloning strategy are available in the supplementary information.

### Generation of vLMα1 transgenic mice and of the vLMα1/APC^+/1638^ mice

Trangenesis is described in details in the supplementary information.

### Azoxymethan (AOM) and AOM/ Dextran Sulfate Sodium (DSS) treatment

Eight week old wildtype (WT) mice and vLMα1 littermates were injected intra-peritoneally (i.p.) with AOM (10 mg/kg, Sigma Aldrich, Lyon, France) once per week for 5 weeks. Animals were sacrificed 9 months after the last AOM injection. For a combined AOM/DSS treatment eight week old WT and vLMα1 littermates were injected i.p. with a single dose of AOM. The day after, 3 % DSS (molecular weight 36,000-50,000 kDa, MP Biomedicals, Illkirch, France) was provided in the drinking water for 5 days. Afterwards mice obtained regular water for 2 months before sacrifice. Tumor size was measured with a caliper and tumor volume was determined using the following calculation V= (width)^2^ × length/2. Colon tumor tissues were prepared, immediately snap frozen for RNA and protein extraction in liquid nitrogen, or were embedded into Tissue-Tek (Labonord, Templemars, France) for immunofluorescence analysis. Samples were stored at −80°C. For conventional histology, samples were fixed in formaldehyde (4%), processed for 10 μm paraffin wax sections and stained with hematoxylin-eosin.

### Cell lines

Control HT29 and HT29LMα1 cells (B8T and H11 clones, respectively) were cultured as previously described (21). Primary cancer associated fibroblasts (CAFs) (52) were infected with a pBABE retroviral vector (Cell Biolabs, Bio-Connect Bv, Huissen, The Netherlands) expressing the hTERT open reading frame, and a pool was selected. The replicative life span of hTERT transduced pool was examined and compared with that of a mock-transduced pool. Immortalized CAFs, mouse embryonic fibroblasts (MEFs kindly provided by R. Chiquet-Ehrismann, Basel, Switzerland) and human normal lung IMR-90 fibroblasts (CCL-186, ATCC, France) were cultured in DMEM supplemented with 10% fetal calf serum (FCS), 1% penicillin-streptomycin (Invitrogen, Life Technologies, Saint Aubin, France). Human immortalized dermal microvascular endothelial cells (HMEC, a gift from Dr E. Van Obberghen-Schilling, Nice, France) were maintained in MCDB 131 medium (Invitrogen) supplemented with 12.5% fetal calf serum, 10 mM glutamine (Invitrogen), EGF (10 ng/mL), bFGF (10 ng/mL), heparin (10 μg/mL) and hydrocortisone (1 μg/mL), all compounds being from Sigma Aldrich, Lyon, France. The human umbilical vein endothelial cells (HUVEC) were grown according to the manufacturer’s instructions in ECGM medium (Promocell, Heidelberg, Germany). Human brain vascular pericytes (ScienCell, USA) were maintained in pericyte medium comprising the basal medium supplemented with 2% fetal bovine serum, 1% penicillin-streptomycin and 1% pericyte growth supplement (ScienCell, USA).

### Generation of LMα1 knock-down cells

Lentiviral strategy is fully detailed in the supplementary information.

### Cell culture assays

CAFs, IMR-90 cells, MEFs, HMEC and pericytes were plated onto uncoated or on LM-111 coated dishes (10μg/cm^2^, L2020, Sigma Aldrich, Lyon, France) and cultured in their appropriate medium. CAFs were then incubated for 24 hours with mouse monoclonal function blocking antibodies against human integrin α2β1 (10μg/mL, BHA2.1, Millipore), integrin β1 (10μg/mL, 4B4, Beckman Coulter, Fullerton, USA) or with rat monoclonal function blocking antibodies against human integrin α6 (10μg/mL, GoH3, Santa Cruz Biotechnology); controls were performed using mouse or rat monoclonal IgG (10μg/mL, Biolegend, Ozyme, St-Quentin-en-Yvelines, France). In certain cases, the AMD3100 inhibitor (10 μg/ml, Merck, Darmstadt, Germany) or the recombinant human CXCL12/SDF1μ protein (100 nM, R & D Systems, Minneapolis, MN, USA) was added to the culture medium for 24 hours concomitantly with antibodies against human integrin α2β1 or integrin β1. Cells were then processed for RNA and protein extraction for subsequent qRT-PCR and Elisa assays.

### Boyden chamber chemo-attraction assay

Chemo-attraction assays were performed in 24 well Boyden chambers with a polycarbonate filter of 8 μm pore size (Falcon, Dutscher, Brumath, France). As a chemoattractant, either conditioned media from HT29 cells (expressing or not LMα1) or purified LM-111 were used. Conditioned media from control HT29 and from HT29LMα1 cells were collected, centrifuged at 900g to remove cell debris and were added to the lower chamber. LM-111 (10μg/cm^2^) was coated on the lower surface of the insert. CAFs, HMEC, HUVEC or pericytes were cultured in the upper chamber (3×10^3^ cells) and incubated for 6 hours at 37°C in 5% CO_2_. Transmigrated cells were then fixed with 4% PFA, stained with DAPI and quantified using the ImageJ software and the analyze particles module (National Institutes of Health, USA).

### *In vitro* angiogenesis assay

The assay is based on a described protocol where endothelial cells are plated on a polymerised gel of basement membrane molecules (Matrigel) on which they form capillary-like structures (53). The basement membrane extract Matrigel (Corning, New York, USA) was loaded (10 μl/well) into 15-well cell culture plates (IBIDI μ-Slide Angiogenesis, Biovalley, Nanterre, France) followed by solidification at 37°C in a humidified incubator for one hour. HUVECs were trypsinized, resuspended at 7 × 10^4^ cells/well, cultured in conditioned medium from CAFs that have been grown previously on plastic or on LM-111 (Sigma, Lyon, France) for 24 hours and treated or not with 1 or 5 μg/ml bevacizumab (Roche, France). After incubation for 7 hours at 37°C, bright field mosaic pictures of the entire well surface were taken (Zeiss Imager Z2 inverted microscope and AxioVision software, Carl Zeiss, Le Pecq, France) at 5X magnification (with a total of 9 pictures per condition). Formation of tubular-like networks was quantified by using the ZEN Blue software (Carl Zeiss, Le Pecq, France) with total number of closed loop structures as read out. A minimum of three independent experiments were performed with five replicates per experiment.

### Cell cycle and apoptosis analysis

**For cell cycle analysis, HT29 control and HT29LMα1 cells** were cultured in 24-well plates (1×10^4^ cells/well) for three days and then treated for 24 hours with recombinant VEGFA165 (10ng/μL, R&D systems, Minneapolis, USA) or VEGFA121 (10ngμL, Prospecbio, East Brunswick, USA). For quantification of apoptosis, cells were treated for 4 hours with the apoptosis inducing agent staurosporine (400nm, Sigma, Lyon, France) as previously described (Qiao et al., 1996), and subsequently treated for 24 hours with the corresponding VEGFA isoforms. After collection, cells were resuspended in 300 μL hypotonic fluorochrome solution (5 μg propidium iodide, 3.4 mmol/L sodium citrate, and 0.1% Triton X-100 in PBS). DNA content was analyzed by a fluorescence activated cell sorter (FACS, Becton Dickinson, San Diego, USA). Ten thousand events per sample were acquired, and cell cycle distribution was determined using the ModFit software. The subG1 apoptotic cell population was quantified by the CellQuest computer software. Apoptosis in tumor samples were measured on protein lysates using the caspase-3/7 colorimetric protease assay kit following the manufacturer’s instructions (caspase-3/7 Colorimetric Protease Assay Kit, Invitrogen, Life Technologies, Saint Aubin, France).

### Surface Plasmon Resonance

Surface Plasmon Resonance-Binding experiments are described in details in the supplementay information.

### Tumor xenograft experiments

Ten million cells of each HT29 cell line (control, HT29LMα1) or 4 million cells of HCT116 (control and HCT116shLMα1) were injected subcutaneously into 8 week old nude MRF1 female mice (Janvier, La Plaine Saint Denis, France). Mice were sacrificed 4 weeks post injection. For histology and immunostaining, tumors were either fixed overnight in 4% PFA and embedded in paraffin or directly frozen in Tissue-Tek on dry ice. For RNA or protein extraction, samples were directly frozen in liquid nitrogen.

### Protein extraction, immunoblotting and ELISA

Proteins were extracted from cells and tissues using lysis buffer (50mM Tris pH 7, 150 mM NaCl, 1% NP-40) and 1% protease inhibitors (cOmplete protease inhibitor cocktail, Roche, Meylan, France). 50 μg of protein lysates (quantified by Bradford assay) were separated by SDS PAGE (6%) and transferred onto nitrocellulose membrane (Millipore, Molsheim, France). Membranes were incubated successively with primary (see **Supplemental Table S7**) and with HRP-coupled secondary antibodies followed by detection upon incubation with ECL (Amersham, GE Healthcare, Velizy-Villacoublay, France). Murine and human specific Quantikine ELISA kits (R&D systems, Minneapolis, USA) were used to determine the amounts of VEGFA and of CXCL12. The CXCR4 Elisa kit was used to detect human CXCR4 (CSB-E12825h; Cusabio, CliniSciences, Nanterre, France). Quantification was performed either by using total tumor or cell protein lysates or using conditioned medium from CAFs, MEFs, IMR-90, HMEC and pericytes, following the manufacturer’s instructions. Absorbance was measured at 450 nm (Biotek plate reader E800, Biotek, Colmar, France).

### Immunohistochemistry and immunofluorescence analysis

For histological analysis, 7 μm paraffin sections were deparaffinized with toluene and stained with periodic acid-Schiff reagent and hematoxylin. The primary antibodies used are listed in the **Supplemental Table S7.** For immunohistochemistry, tissue sections were deparaffinized with toluene, then boiled with the antigen retrieval sodium citrate buffer (pH 6) for 10 min. Sections were incubated with primary antibodies overnight at 4°C. Slides were thereafter incubated with biotinylated secondary antibodies (Vector Laboratories, Eurobio/Abcys, Les Ulis, France), amplified with the ABC Elite Vectorstain kit and developed with the DAB kit from Vector Laboratories. Slides were examined using the Zeiss Axio Imager A1 microscope equipped with an A-Plan x5/0.12, an A-Plan x20/0.45 objective and a Zeiss Axiocam Icc3 color camera (Carl Zeiss, Le Pecq, France). For immunofluorescence staining, 7 μm cryosections were incubated overnight with primary antibodies, washed three times in PBS and incubated for one hour with Alexa 488- or cyanine 3-conjugated secondary antibodies (Jackson ImmunoResearch Laboratories, West Grove, PA). After washing, nuclei were stained with DAPI (1/30000) and embedded using the FluorSave reagent (Calbiochem-Merck, Lyon France). Slides were examined using an epifluorescence Zeiss Axio imager 2 microscope equipped with a Plan Apochromat x20/0.8, a Plan Apochromat x40/0.95 objectives and an apotome module. Pictures were taken with a Zeiss Axiocam MRm black and white digital camera. All images were acquired using the Zeiss Axiovision software. Control sections were processed as above with omission of the primary antibodies. Quantifications of immunofluorescence and immunohistochemistry surface signals were done using the ImageJ software and the analyze particles module (National Institutes of Health, USA). Several images per tumor were taken using a 20x objective to cover most of the tumor surface. Data are presented as average area fraction per tumor in all defined groups. Quantification of the pericyte coverage index of vessels was defined by dividing signals for the average area fraction of NG2 by the average area fraction of CD31.

### Gene expression analysis

Gene expression analysis and experiments are described in the supplementary information.

### Proximity ligation assay

The proximity ligation assay was employed following the manufacturer’s recommendations (Duolink In situ Orange starter kit mouse/rabbit; Sigma-Aldrich, France). CAFs were seeded (2×10^4^ cells/well) on uncoated or LM-111 (10μg/cm^2^; Sigma-Aldrich, France) coated Lab-Tek II chamber slides (Dutscher). After cell adhesion and spreading, CAFs were starved overnight, fixed with 1% PFA in PBS for 10 min and permeabilized with 0.1% Triton-PBS for 10 min. Then cells were incubated overnight at 4°C with both mouse anti-integrin α2β1 (BHA2.1 clone; 1/100; Millipore) and rabbit anti-CXCR4 (Ab2074; 1/100; Abcam) antibodies. Ligation, amplification and detection of integrin α2β1 and CXCR4 interactions were visualized using the Duolink kit following manufacturer’s instructions. In this procedure, bright fluorescent dots were observed when the two molecules are in close proximity. Images were taken using the epifluorescence Zeiss Axio imager 2 microscope and quantification of dots was performed using Image J software on 11 random fields obtained from 2 independent experiments.

### Statistical analysis

Statistical significance of results was analyzed using the GraphPad Prism program version 5.04 and the R open source software version 3.2.1. The Shapiro-Wilk normality test was used to confirm the normality of the data. The statistical difference of Gaussian data sets was analyzed using the Student unpaired two-tailed t test, with Welch's correction in case of unequal variances and the one way ANOVA test followed by a Tukey's multiple comparison post-test was used for multiple data comparison. For data not following a Gaussian distribution, the permutation test was used and the one-way ANOVA test followed by the permutation multiple comparisons post-test was used for multiple data comparison. Illustrations of these statistical analyses are displayed as the mean +/− standard deviation (SD). Contingency was analyzed using the chi-square test. p-values smaller than 0.05 were considered as significant. *, p<0.05, **, p < 0.01, ***, p < 0.001, ****, p < 0.0001.

## Abbreviations

BM: basement membrane
ECM: extracellular matrix
LM: laminin
vLMα1: villin-LMα1 transgenic mice
CAFs: cancer-associated fibroblasts
IF: immunofluorescence
VEGF: vascular endothelial growth factor.

## Additional information

## Financial support

This work was supported by INSERM, Ligue contre le Cancer (OL, GO), Institut National du Cancer (PSA, GO), the Association pour la Recherche sur le Cancer and l’Hôpital de Hautepierre, Strasbourg (GO). EMB, TR and CS were recipients of a fellowship from the Ligue contre le Cancer and IJ was supported by the Fondation pour la Recherche Médicale.

## Conflicts of interest

The authors declare no conflicts of interest.

## Supplemental information

Supplemental Information includes supplemental materials and methods, Six supplemental figures, eight supplemental tables, the legends to the supplemental display items and references.

## Acknowledgments

We thank R. Fodde (Department of Human Genetics, Leiden University Medical Center, The Netherlands) for APC^+/1638N^ mice and the Mouse Clinical Institute (Illkirch-Graffenstaden, France) for the generation of transgenic mice. We also thank R. Chiquet-Ehrismann (Friedrich Miescher Institute for Biomedical Research, Basel, Switzerland) for Mouse Embryonic Fibroblasts, E. Van Obberghen-Schilling (University of Nice-Sophia Antipolis, Nice, France) for HMECs, P. Yurchenco (Robert Wood Johnson Medical School, Piscataway, NJ, USA) for mouse LMα1 cDNA, L. Sorokin (Institute for Physiological Chemistry and Pathobiochemistry, University of Münster, Münster, Germany), H. Kleinman (The George Washington School of Medicine, Washington, DC, USA), T. Sasaki (Shriners Hospitals for Children, Portland, OR, USA) and D. Gullberg (Department of Biomedicine, University of Bergen, Bergen, Norway) for antibodies and, the Centre de Ressources Biologiques (Hôpital de Hautepierre, Strasbourg, France) for the human tissue biopsies. Sequencing and microarray were performed by the IGBMC Microarray and Sequencing platform (http://genomeast.igbmc.fr).

